# A high-quality, chromosome-level genome assembly of the Black Soldier Fly (*Hermetia Illucens* L.)

**DOI:** 10.1101/2020.11.13.381889

**Authors:** Tomas N. Generalovic, Shane A. McCarthy, Ian A. Warren, Jonathan M.D. Wood, James Torrance, Ying Sims, Michael Quail, Kerstin Howe, Miha Pipan, Richard Durbin, Chris D. Jiggins

## Abstract

**Background:** *Hermetia illucens* L. (Diptera: Stratiomyidae), the Black Soldier Fly (BSF) is an increasingly important mass reared entomological resource for bioconversion of organic material into animal feed.

**Results:** We generated a high-quality chromosome-scale genome assembly of the BSF using Pacific Bioscience, 10X Genomics linked read and high-throughput chromosome conformation capture sequencing technology. Scaffolding the final assembly with Hi-C data produced a highly contiguous 1.01 Gb genome with 99.75% of scaffolds assembled into pseudo-chromosomes representing seven chromosomes with 16.01 Mb contig and 180.46 Mb scaffold N50 values. The highly complete genome obtained a BUSCO completeness of 98.6%. We masked 67.32% of the genome as repetitive sequences and annotated a total of 17,664 protein-coding genes using the BRAKER2 pipeline. We analysed an established lab population to investigate the genomic variation and architecture of the BSF revealing six autosomes and the identification of an X chromosome. Additionally, we estimated the inbreeding coefficient (1.9%) of a lab population by assessing runs of homozygosity. This revealed a plethora of inbreeding events including recent long runs of homozygosity on chromosome five.

**Conclusions:** Release of this novel chromosome-scale BSF genome assembly will provide an improved platform for further genomic studies and functional characterisation of candidate regions of artificial selection. This reference sequence will provide an essential tool for future genetic modifications, functional and population genomics.

## Background

The Black Soldier Fly (BSF; Figure 1), *Hermetia illucens*, Linnaeus, 1758 (Diptera: Stratiomyidae; NCBI: txid343691) is a species of growing interest in both entomophagy and bioremediation. Endemic to tropical and sub-tropical regions of the Americas, BSF are now distributed globally extending to temperate regions of Europe and Asia through commercialisation and human mediated expansion [1–3]. The increasing popularity of BSF in insect farming is due to the high feed-to-protein bioconversion rates of BSF larvae coupled with a generalist diet. Conversion efficiency of BSF larva is higher than other traditional edible insects such as *Tenebrio molitor* (Yellow mealworm) [4]. Due to the high protein (40%) and lipid (35%) content of BSF larvae, they are now a European Union approved feed ingredient in aquaculture and poultry farms as a replacement to inefficient and unsustainable fish and soybean meal [5,6]. Additionally, the rich biomass of BSF larvae have led to the resource exploitation of lipids and chitin for the cosmetic industry, as a source of biofuel production and recently shown promise as a source of antimicrobials [7–10]. With short generation times, large brood sizes (~900 eggs per clutch) and voracious feeding behaviour, the BSF is the optimal species for insect mass rearing [11,12].

**Figure 1.**
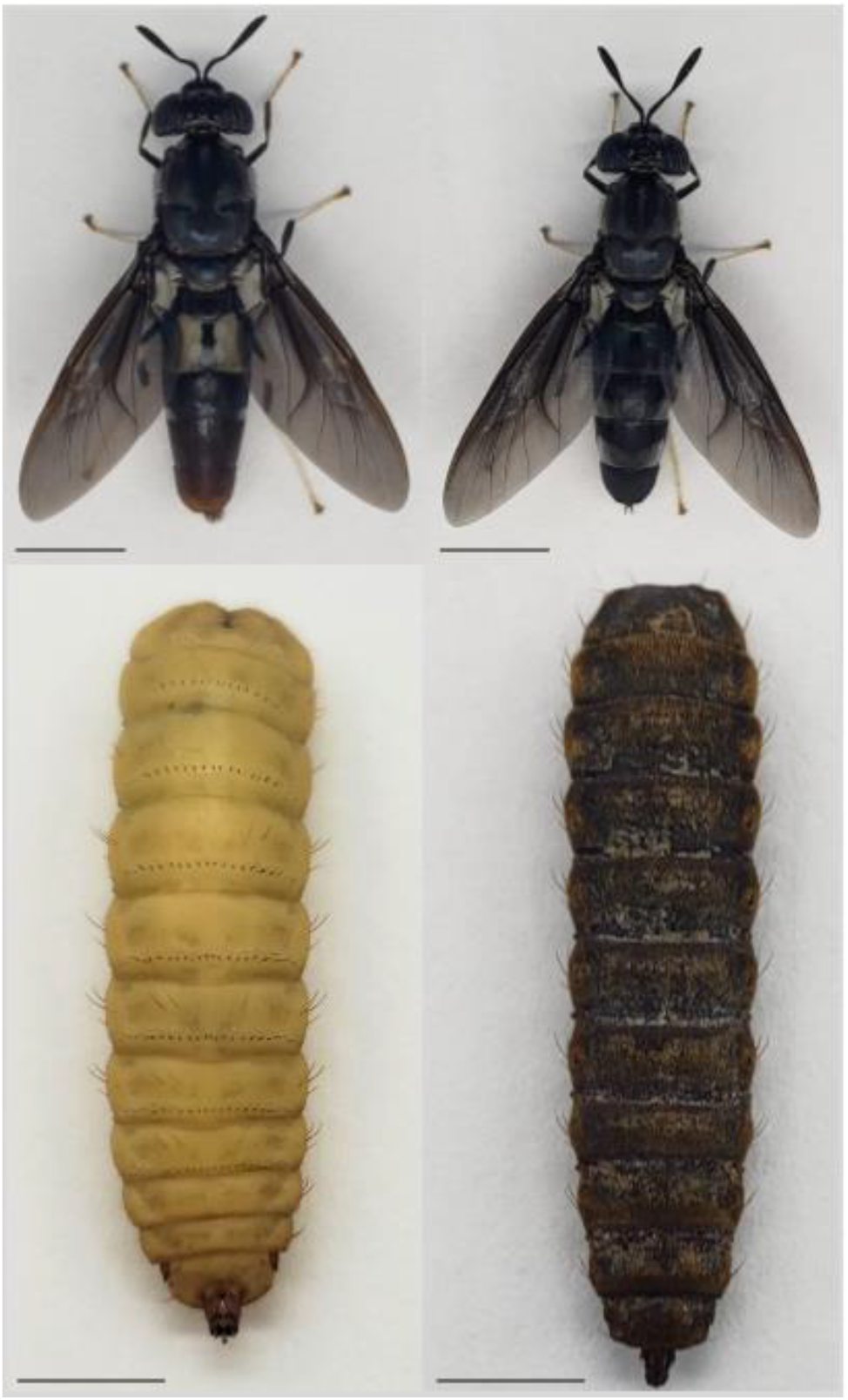
*Hermetia illucens* key life stages. Dorsal view of *Hermetia illucens* adult male (upper left), adult female (upper right), larvae (5^th^ instar; lower left) and pupa (lower right). Adult sex identified by genital shape and posterior colour. Scale bar = 5 mm. Photos: T.N.G.

Global food security and waste management systems are increasingly under threat from a growing human population. An increase in food production to feed a population of over 9 billion by 2050 will require a demand for protein in excess of 270 million tonnes [13,14]. With one third of this produce likely to be processed as food waste, a transition to a more sustainable agricultural model is essential [14]. Shifting to a circular bioeconomy utilising insect biomass can lead to a more sustainable global food industry [15]. With high nutrient content and the ability to upcycle organic waste streams, the BSF is the most exploited species in the growing insect farming industry [16]. However, whilst most research aims to optimise biomass yield, there remains a lack of understanding surrounding the genetics of the BSF.

Whilst genomic resources within the Diptera order are well established through databases such as FlyBase, the resources available for BSF are limited [17]. A reference genome of the BSF has recently been released but is relatively fragmented [18]. The expanding industry of BSF mini-livestock farming must rapidly match the high genomic standards of other agricultural animals if it is to become a successful and well - established practice [19]. It is essential to characterise genetic traits and beneficial phenotypes to provide longevity to this novel agriculture market. Recent advancements in sequencing technology and high-throughput projects including the Darwin Tree of Life has enabled the assembly of many non-model reference genomes [20,21]. Arthropod genome assembly projects can be hindered by limited starting material, ploidy level, repeat rich genomes and high heterozygosity [22]. Resolving genome heterozygosity remains a major challenge in diploid and polyploid assembly projects [23]. Nonetheless, integrating long read and linked-read sequencing has greatly facilitated *de novo* assemblies [24]. Combined with high-throughput chromosome conformation capture (Hi-C) sequencing, the assembly of chromosome-level reference genomes in vertebrates, invertebrates and plants is far more approachable [25–27].

Here we present a chromosome-level, 1.01 Gb genome assembly for the economically important insect, *H. illucens* (BSF). We demonstrate the combination of Pacific Bioscience (PacBio) long read, 10x Genomics Linked read and chromosomal conformation capture sequencing data in assembling a highly contiguous, complete and accurate genome. We identify and mask novel repeat elements to annotate the BSF genome using available transcriptome evidence to provide an essential resource for currently understudied BSF genetics. We also use genome re-sequence data to identify an active sex chromosome, assess the genomic variation and the level of inbreeding in an established lab population. As the first chromosome-scale assembly available for the BSF this resource will enable the genetic characterisation and understanding of unique traits and will further the development of genetic studies including population genetics and genetic manipulation.

### Insect husbandry and collection

A captive population of *Hermetia illucens* was supplied by Better Origin (Entomics Biosystems Limited) and reared under controlled conditions at the University of Cambridge, Zoology department (Cambridge, UK). A mating pair were isolated, and the offspring reared under conditions of 29 ± 0.8 °C, 60% relative humidity on a 16:8-hour light: dark cycle. Larval offspring were fed twice weekly ad libitum on Better Origin (Cambridge, UK) ‘BSF Opti-Feed’ mixed 30:70 with non-sterile H_2_O under the same conditions. Pre-pupae were transferred to medium-grade vermiculite (Sinclair Pro, UK) for pupation and emerged adults moved to a breeding cage (47.5 × 47.5 × 93 cm; 160 μm aperture). An adult female and male of the founding population and two offspring pupae were collected and stored at −80°C until processed for sequencing. Additional offspring from the same pair were used to establish a BSF colony line named “EVE” for future analysis. A sample (n=12) of the EVE colony from the eighth-generation post - lab establishment were also collected and stored at −80°C until processed for sequencing.

### Genome size and Heterozygosity estimates

Genomic DNA (gDNA) was extracted from the thorax of the mature adults using the Blood & Cell Culture DNA Midi Kit (Qiagen, Germany) following the manufacturers protocol. Paired-End (PE) libraries were produced and sequenced on the Illumina HiSeq × Ten platform (Illumina, United States) at the Wellcome Sanger Institute (Cambridge, UK). Sequencing of the adult female and male produced 150.38 Gb and 162.42 Gb respectively (Table S1). Sequencing data was used to estimate genome size, heterozygosity and genomic repeats using GenomeScope [28]. Distribution of k-mers in both individuals using k = 31 produced a diploid peak set. We estimated a genome size of 1.06 Gb, provisional repeat content of 46.25% and mean heterozygosity of 1.81% (Figure S1 & Table S2).

### Genome library construction & sequencing

An offspring pupa from the isolated pair was collected to prepare libraries for both Pacific Biosciences (United States) and 10X Genomics Chromium linked-read (10X Genomics, United states) sequencing (Table S1). All *de novo* genome sequencing was carried out at the Wellcome Sanger Institute (Cambridge, UK). High-molecular weight DNA was extracted from the pupal offspring using a modified MagAttract HMW DNA protocol (Qiagen, Germany) and a PacBio SMRT sequencing library was prepared. Eight Single Molecule Real Time (SMRT) cells (1M v2) using version 2.1 chemistry of the PacBio Sequel platform generated 75.17 Gb subreads with an N50 of 22.71 kb. An additional sequencing library was constructed and sequenced on a single 8M SMRT cell on the Sequel II platform using version 0.9 sequencing chemistry to produce an additional 105.69 Gb of data with an N50 of 14.58 kb from the same individual, giving a total of 180.86 Gb. A 10X Genomics Chromium linked read 150 bp PE library was also prepared from the gDNA of the same individual. Sequencing of the linked read library on the Illumina HiSeq × Ten platform (Illumina, United States) produced 149.78 Gb of raw data. A Hi-C PE library was constructed from the tissue of a sibling pupa and 150 bp PE reads were sequenced on the Illumina HiSeq × Ten platform. Hi-C sequencing produced 144.5 Gb of raw data.

### Genome assembly

Due to the high predicted heterozygosity of the BSF genome we employed FALCON-Unzip, a *de novo* diploid-aware assembler of PacBio SMRT sequence data prior to scaffolding [29]. The FALCON-Unzip algorithm utilises the hierarchical genome assembly process (HGAP) enabling haplotype resolution. Raw PacBio data containing reads longer than 5 kb were selected for error correction and consensus calling. The intermediate BSF genome assembled into a genome size of 1.09 Gb into 140 contigs with an N50 value of 13.7 Mb. The FALCON-Unzip draft assembly was purged of duplicate sequences using purge_dups v0.0.3 [30]. The 10X linked reads were used to scaffold the draft FALCON-Unzip assembly using scaff10x [31]. The assembly was polished with one round of arrow [32] using the error corrected reads of FALCON-Unzip. The Hi-C data was mapped to the intermediate assembly in a second round of scaffolding using BWA [33]. A Hi-C contact map was generated and visualised in HiGlass [34]. This final draft assembly was manually curated to remove contaminants, correct structural integrity and assemble and identify chromosome-level scaffolds using gEVAL [35,36]. Remaining haplotype duplication was purged manually into an alternative haplotype genome. The final Hi-C contact map (Figure 2) was visualised in HiGlass [34]. The resulting chromosome-level assembly deemed “iHerIll” consisted of a total length of 1,001 Mb, contig N50 of 16.01 Mb and a scaffold N50 of 180.36 Mb (Table 1; Table S3). We anchored 99.75% of assembled scaffolds into seven chromosomes leaving 13 unplaced scaffolds (2.53 Mb; 0.25%; Table S4).

**Table 1.** Summary statistics of *Hermetia illucens* and selection of Diptera genomes. Assembly statistics for Diptera genomes generated using assembly-stats script on the iated reference genome. BUSCO score generated from the ‘insecta_odb9’ database (n = 1658). BUSCO statistics C: complete, S: single-copy, D: duplicated, F: fragmented, issing. * denotes contig number where the assembly contains no scaffolds.

| Species name | Genome size | Scaffold number | Contig N50 | Scaffold N50 | Gaps | N count | GC (%) | BUSCO (%) |  |  |  |  |
| --- | --- | --- | --- | --- | --- | --- | --- | --- | --- | --- | --- | --- |
|  |  |  |  |  |  |  |  | C | S | D | F | M |
| *Hermetia illucens* (iHerIII) | 1.01 Gb | 20 | 16.01 Mb | 180.36 Mb | 112 | 26,439 | 42.47 | 98.60 | 97.80 | 0.80 | 0.50 | 0.90 |
| *Hermetia illucens*<br>(GCA_009835165.1) | 1.10 Gb | 2,806 | 231.07 kb | 1.70 Mb | 24,796 | 14,305,322 | 42.46 | 98.90 | 91.10 | 7.80 | 0.60 | 0.50 |
| *Drosophila melanogaster*<br>(GCA_000001215.4) | 143.73 Mb | 1,870 | 21.49 Mb | 25.29 Mb | 572 | 1,152,978 | 41.67 | 99.70 | 99.00 | 0.70 | 0.20 | 0.10 |
| *Drosophila virilis*<br>(GCA_000005245.1) | 169.7 Mb | 13,530 | 5.10 Mb | 31.08 Mb | 166 | 16,600 | 40.43 | 99.10 | 98.10 | 1.00 | 0.40 | 0.50 |
| *Musca domestica*<br>(GCA_000371365.1) | 750.40 Mb | 20,487 | 11.81 kb | 226.57 kb | 102,610 | 58,683,792 | 32.37 | 98.60 | 96.90 | 1.70 | 0.40 | 1.00 |
| *Stomoxys calcitrans*<br>(GCA_001015335.1) | 971.19 Mb | 12,042 | 11.31 kb | 504.65 kb | 129,113 | 150,529,258 | 40.30 | 98.40 | 97.70 | 0.70 | 1.00 | 0.60 |
| *Glossina morsitans morsitans*<br>(GCA_001077435.1) | 363.11 Mb | 24,071* | 49.77 kb | NA | 0 | 0 | 34.12 | 98.90 | 96.60 | 2.30 | 0.60 | 0.50 |
| *Aedes aegypti*<br>(GCA_002204515.1) | 1.28 Gb | 2,310 | 11.76 Mb | 409.78 kb | 229 | 22,935 | 38.18 | 98.90 | 94.50 | 4.40 | 0.40 | 0.70 |
| *Culex quinquefasciatus*<br>(GCA_000209185.1) | 579.04 Mb | 3,171 | 28.55 kb | 486.76 kb | 45,500 | 39,082,744 | 34.89 | 96.70 | 91.80 | 4.90 | 0.80 | 2.50 |

**Figure 2.**
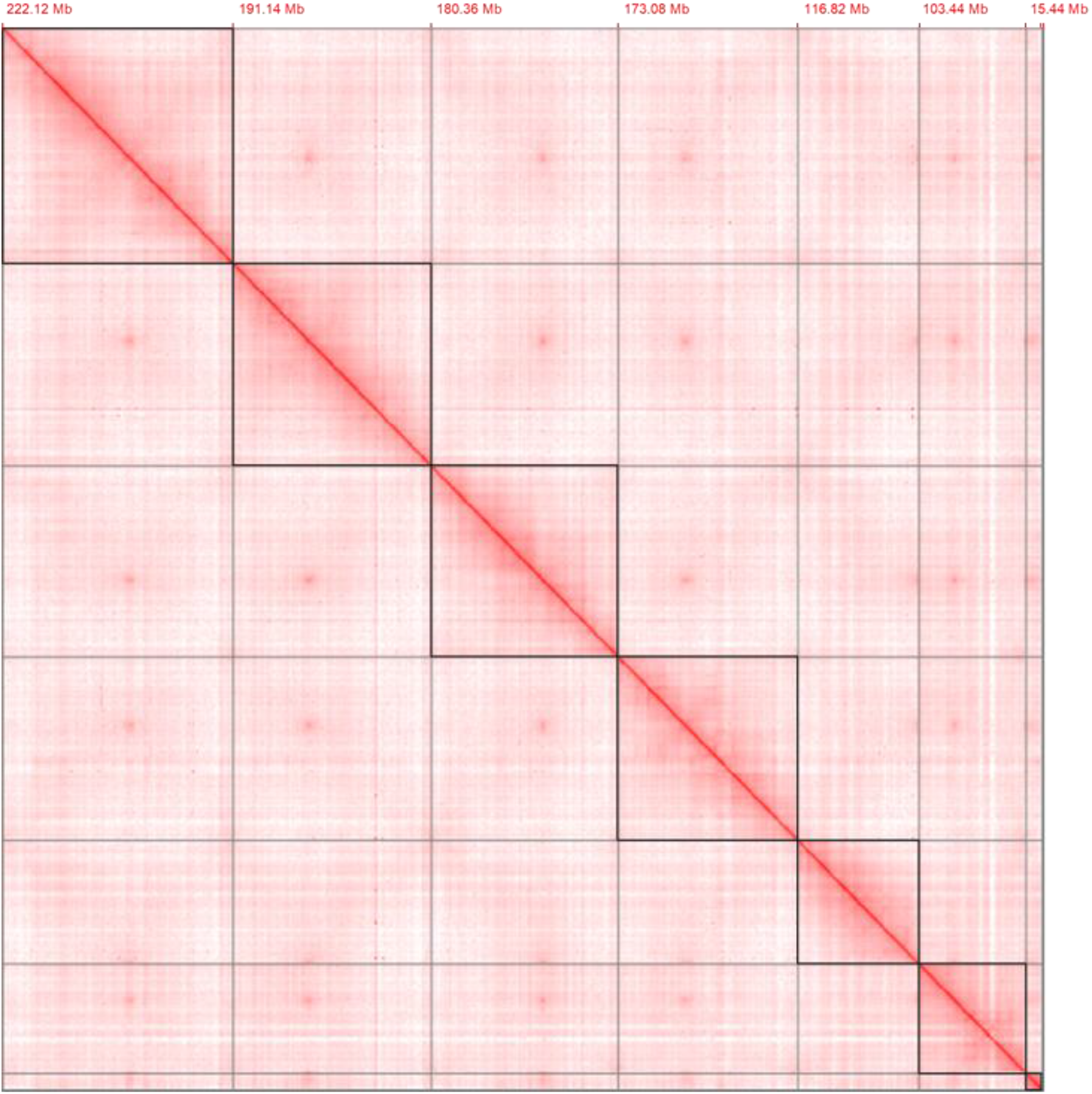
Curated Hi-C contact map of chromosomal interactions. Super-scaffold chromosomes (n=7) are highlighted within black frames and annotated with assembled length at the beginning of each chromosome (interactive map available on https://higlass-grit.sanger.ac.uk/l/?d=OjQNRJcmTgKyKfvkk3yODg).

### Genome quality evaluation

To assess the quality of the reported assembly we evaluated both genome completeness and contamination. We used BUSCO v3.0.2 (Benchmarking Universal Single-Copy Orthologs) [37] to identify conserved genes within the ‘insecta_odb9’ and ‘diptera_odb9’ databases. The final assembly covered 98.60% and 92.60% of the Insecta and Diptera BUSCO core genes respectively. Within the Insecta BUSCO completeness score of the reported genome 1,622 (97.80%) were single copy with 15 (0.9%) missing (Figure 3; Table S5). Contamination was assessed using the BlobToolKit environment v1.0 [38] to screen for contaminant sequence. BlobToolKit identified 99.89% of the raw PacBio data assembled exclusively arthropod sequence (Figure S2). Whilst Arthropoda was the highest abundant identified phyla, 1,137,222 bp (0.11%) obtained no significant taxonomic identification. Therefore, our assembly does not include any significant assembled contaminate sequences providing low likelihood of off-target mapping effects for future studies. Whereas the majority of the genome sequence was identified as Diptera (77.83%), segments of closest sequence similarity to Siphonaptera (11.63%), Hymenoptera (10.29%), Coleoptera (0.09%) and Lepidoptera (0.05%) were also identified.

**Figure 3.**
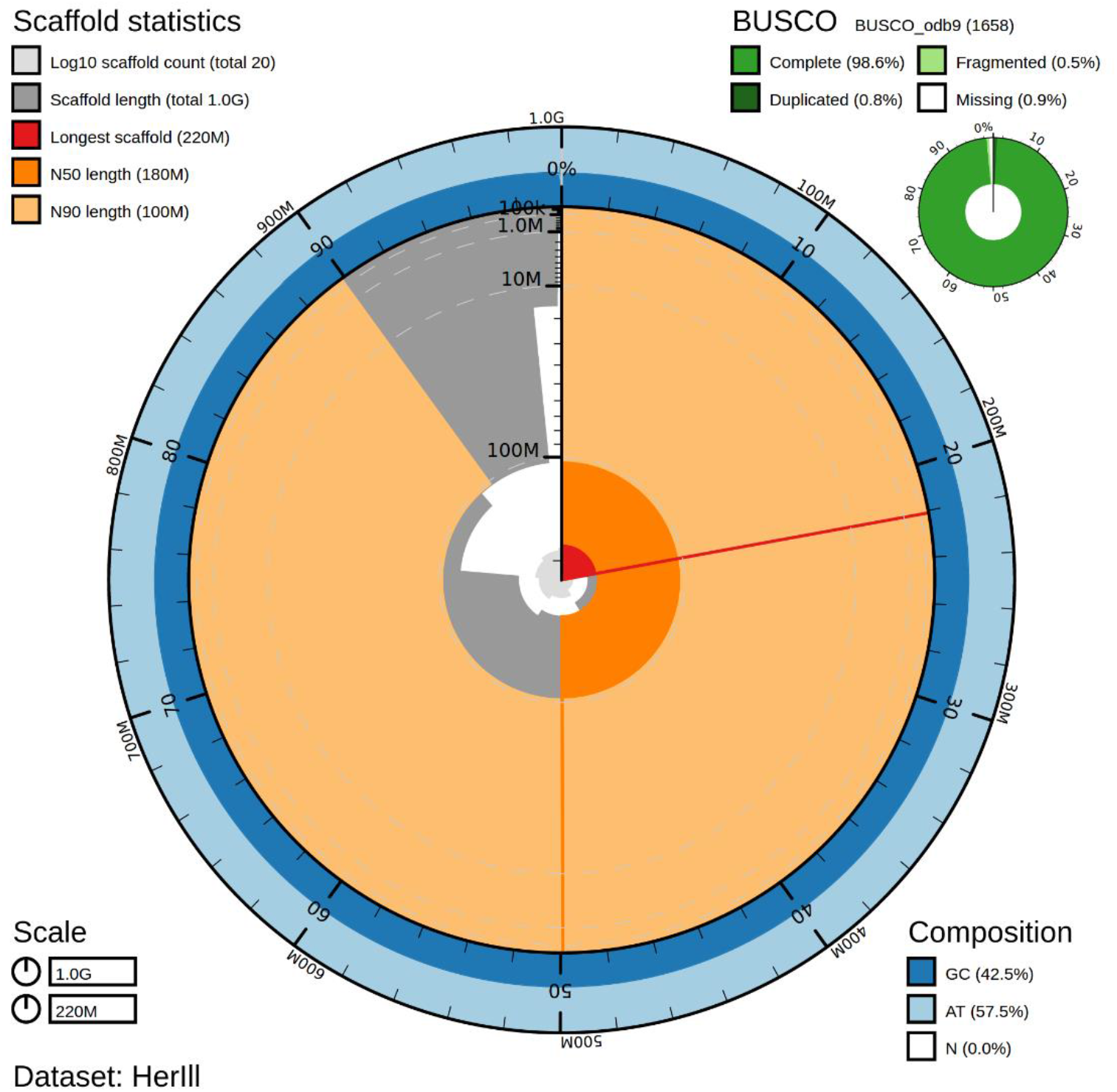
BlobToolKit snail plot of the *Hermetia illucens* assembly. Genome assembly statistics of iHerIll visualised as a snail plot containing BUSCO ‘insecta_odb9’ completeness scores, scaffold assembly statistics and sequence composition proportions.

### Repeat sequence identification

To quantify the repetitive regions within the BSF genome we modelled and masked a *de novo* library of repetitive sequences using RepeatModeler v2.0.1 and RepeatMasker v4.0.9 [39]. A custom repeat consensus database was built, and repeat elements classified using RepeatModeler “-engine ncbi”. The Dfam_Consensus-20181026 [40] and RepBase-20181026 [41] databases were combined with the custom repeat database. Using the merged database RepeatMasker was used to identify and soft mask repetitive regions using the RMBlast v2.6.0 sequence search engine. Repeatmasking resulted in a total of 67.32% (676,593,256 bp) of the genome being identified as repeat regions (Table 2). We identified Long Interspersed Nuclear Elements (LINEs; 44.85%) as the most abundant class of repetitive elements followed by a high proportion of unclassified repeats (13.81%). This repeat analysis identifies comparable statistics to the identified repeats of the previously published BSF genome (65.76%).

**Table 2.** Genome annotation and repeat statistics of the *Hermetia illucens*.

| Genome annotation statistics |  | *Hermetia illucens* assembly |  |  |
| --- | --- | --- | --- | --- |
|  |  |  | # |  |
| Protein-coding genes |  |  | 17,664 |  |
| BUSCO statistics for protein-coding gene annotation |  | % | # |  |
| Insecta<br>(n:1658) | Completeness | 98.20 | 1,627 |  |
|  | *Complete Single-copy* | 91.10 | 1,510 |  |
|  | *Complete Duplicated* | 7.10 | 117 |  |
|  | Fragmented | 0.70 | 12 |  |
|  | Missing | 1.10 | 19 |  |
| Diptera<br>(n:2799) | Completeness | 95.60 | 2,674 |  |
|  | *Complete Single-copy* | 86.10 | 2,409 |  |
|  | *Complete Duplicated* | 9.50 | 265 |  |
|  | Fragmented | 2.50 | 70 |  |
|  | Missing | 1.90 | 55 |  |
| Repeat statistics |  | bp | % | # |
| DNA Elements |  | 38,102,124 | 3.79 | 108,133 |
| LTR |  | 45,335,874 | 4.51 | 72,542 |
| LINES |  | 450,717,046 | 44.85 | 946,086 |
| SINES |  | 3,620,717 | 0.36 | 16,138 |
| Unclassified |  | 138,817,495 | 13.81 | 485,192 |
| Total interspersed repeats |  | 676,593,256 | 67.32 | 1,628,091 |

### Genome annotation

Genome annotation of the assembly was produced using the BRAKER2 pipeline v2.1.5 [42–49]. Raw RNA-seq reads were obtained from whole larva (study accession: PRJEB19091) [50] and a separate study using the full BSF life cycle; egg (12 & 72 hours-old), larva (4, 8 & 12 days-old), pre-pupa and pupa (both early and late stages) including both male and female adults (BioProject ID: PRJNA573413) [18]. Arthropod proteins were obtained from OrthoDB [51]. RNA-seq reads were filtered for adapter contamination and low-quality reads using Trimmomatic v0.39 [52] followed by quality control pre and post trimming with fastqc v0.11.8 [53]. RNA-seq reads were mapped to the assembly using STAR (Spliced Transcripts Alignment to a Reference) v2.7.1 [54] in 2-pass mode. Protein hints were prepared as part of the BRAKER2 pipeline using ProtHint v2.5.0. BRAKER2 *ab initio* gene predictions were carried out using homologous protein and de novo RNA-seq evidence using Augustus v3.3.3 [42] and GeneMark-ET v4.38 [45]. Genome annotations were assessed for completeness using BUSCO v3.0.2 (--m prot) ‘insecta_odb9’ and ‘diptera_odb9’ databases [37]. The BRAKER2 pipeline annotated 17,664 protein-coding genes which provided BUSCO completeness scores of 98.2% and 95.6% for Insecta and Diptera core gene datasets respectively (Table 2).

### Comparative genome assembly analysis

We evaluated the assembly statistics of the iHerIll genome for completeness, contiguity and correctness with related Diptera species and the only published BSF reference, GCA_009835165.1 [18]. Assembly statistics (Table 1) for publicly available Diptera genomes were generated using assembly-stats [55]. To assess the completeness of the iHerIll assembly we employed BUSCO v3.0.2 ‘insecta_odb9’ and ‘diptera_odb9’ core gene sets and compared results to related Diptera. The BUSCO results were comparable between existing high-quality Diptera genomes and iHerIll. In comparison with the previous BSF GCA_009835165.1 assembly there were just 15 BUSCO missing from the iHerIll assembly whereas seven were missing in the GCA_009835165.1 assembly (Table S5). We assemble a high proportion of single copy orthologs (97.8%) with little gene duplication (0.8%) within the iHerIll assembly indicating that many duplicated genes remain in the GCA_009835165.1 assembly (7.8%), likely due to unresolved heterozygous regions. Our iHerIll assembly is one of the highest quality assembled dipteran genomes available, assembled into the smallest number of scaffolds with the largest scaffold N50 value amongst sampled Diptera (Table 1).

To measure genome contiguity and correctness we incorporated a quality assessment tool, QUAST v5.1.0 [56]. The GCA_009835165.1 assembly was first filtered to purge highly fragmented contigs < 10 kb, retaining 99.86% of the original sequence and aligned to the reported iHerIll assembly. Genome alignment statistics generated using QUAST (“--large -eukaryote --min-alignment 500 --extensive-mis-size 7000 --min-identity 95 --k-mer-stats --k-mer-size 31 --fragmented”) provided NA50 and LA50 metrics based off aligned sequences, enabling comparable results to be drawn between the two assemblies and identify potential misassembly events. We confirm the much higher contiguity of iHerIll compared with GCA_009835165.1 (Figure S3) and identify 787 potentially misassembled contigs in GCA_009835165.1, together with ten contigs that do not align at all to iHerIll (Table S6). We next produced whole genome alignments using mummer v3.23 [57] and visualised alignments using dnanexus [58]. The GCA_009835165.1 assembly showed substantial full-length alignment to iHerIll scaffolds but with several insertions and deletions between the assemblies (Figure S4). To test whether the BSF reference assemblies harboured unique genomic sequence we hard masked both genomes using the custom repeat library for re-alignment. The GCA_009835165.1 assembly contained 302.4 kb of non-repetitive DNA sequence that did not align to the iHerIll assembly. None of the iHerIll genome failed to align to GCA_009835165.1. Severe inbreeding effects, hybridisation and extensive chromosomal rearrangements here may promote high levels of sequence divergence between populations [59]. It is likely that some of the contigs that fail to align properly represent true biological differences rather than misassemblies. Whilst both genomes support complete assemblies, a previous study of the BSF mitochondrial cytochrome c oxidase I (CO1) gene indicates a high level of genetic diversity that is suggestive of a larger species complex than originally thought [60]. Assembly of further high-quality genomes from additional BSF lines will be essential to reveal the extent of genomic diversity within the BSF species-complex.

### Genomic variation

A sample of 12 individuals from generation eight of the sampled BSF line ‘EVE’ were sequenced on the BGI-seq platform. DNA from adult whole thorax tissue was extracted using the Blood & Cell Culture DNA Midi Kit (Qiagen, Germany). Sequencing libraries for 12 individuals of 150 bp PE PCR-free BGI-seq Whole Genome Sequencing (WGS) were prepared and sequenced to an average genome coverage of 25x by BGI (BGI, Hong Kong). Sequencing data was quality checked using FastQC v0.11.5 [61] pre and post adapter trimming with cutadapt v1.8.1 [62]. Raw data was mapped to the assembled genome using BWA v0.7.17 [33], sorted with samtools v1.9 [63] and duplicates removed using picard v2.9.2 [64]. Variant calling was carried out using bcftools mpileup v1.9 [63] and filtered using vcftools v0.1.15 [65]. To obtain high quality Single Nucleotide Polymorphism (SNP) datasets we removed indels (--remove-indels), applied a minimum and maximum mean depth (--min-mean-DP 12; --minDP 12; --max-meanDP 30; --maxDP 30), a minimum quality threshold (--minQ 30) and removed sites missing > 15% of data (--max-missing 0.85). Genome nucleotide diversity (π) and Tajima D were calculated over 20 kb windows using popgenWindows.py (-w 20000 -m 100 -s 20000) [66] and vcftools (--TajimaD 20000) respectively. Further filtering for minor allele frequency (MAF) filter (--maf 0.05) was also applied for runs of homozygosity (ROH) analysis. Runs of homozygosity (ROH) were generated with the detectRUNS v0.98 R package [67] using sliding windows (windowSize=15; threshold=0.05; minSNP=15, minLengthBps = 200000) and default parameters. Inbreeding coefficients were generated using the calculation 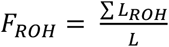 where L is total autosome length.

We estimated mean genome wide nucleotide diversity (π) as 0.017 and Tajima D to be 1.58 (Table S7). Chromosome five exhibited the only negative Tajima D and the lowest nucleotide diversity. We identified a total of 444 genome wide ROH in the sample population using 3,834,541 high-quality SNPs of which 96.4% were < 1 Mbp in length (Table S7). The remaining 3.6% were deemed long ROH at a length ≥ 1 Mbp. Islands of ROH were consistent across individuals within the population (Figure S6). Mean ROH length in the EVE line was identified as 393,812 bp with 43.5% of the total runs located on chromosome five (Table S7; Figure S5). Half of the identified long ROH were located within a 17.6 Mb region on chromosome five containing 239 annotated genes. The genome wide inbreeding coefficient estimated from these ROH was 0.019 (excluding the identified sex chromosome, see below). This established captive EVE line does not appear to be hindered by inbreeding depressions unlike previous inbred populations which have experienced severe colony crash events [68]. However, small sample size (n=12) may provide a bias distribution of allele frequencies for inbreeding calculations. Sequence diversity of this BSF line remains low whilst complimented with a strongly positive Tajima D statistic is consistent with a recent population bottleneck. Reduced nucleotide diversity and Tajima D on chromosome five are consistent with patterns of extended homozygosity, particularly within the 79.9 to 97.5 Mb region of interest (Figure S6). An excess of rare alleles in this region may indicate a potential selective sweep. High proportions of short ROH indicates many recombination events in the life history of this population likely as a product of long-term domestication [69]. However, long ROH indicates recent inbreeding as a result of the founder effect during establishment of the EVE line from an extremely small population [70]. Nonetheless, localised islands of ROH, reduced diversity and Tajima D may also indicate potential candidate regions of adaptation as a product of extensive domestication, with chromosome five of interest for future population genetic investigations. Monitoring strategies using genomic data are essential for conservational genetics and tracking pedigrees in farmed populations [71]. Utilising data such as these will aid in preventing BSF colony collapse in commercial facilities. However, it also introduces the potential for selective breeding programs, enabling the improvement of highly productive life history traits through artificial selection.

### Sex chromosome identification

Further investigation identified a sex chromosome using re-sequence data from both female (n=7) and male (n=5) adults. The final assembly was hard masked of repetitive sequence using the custom repeat library. Sex specific sequence data was mapped using BWA v0.7.17 [33] and merged using samtools v1.9 [63]. Mean depth of coverage was generated over 20 kb genome wide windows in 20 kb steps using samtools v1.9 [63] (depth -aa) including unmapped regions [63], genomics.py and windowscan.py (--writeFailedWindows -w 20000 -s 20000) [66]. Mean depth was plotted as log2 fold change (male/female) and visualised in R v3.3.2 [72] using the ggplot2 package [73]. Chromosome one to six exhibited the coverage of an autosome whilst chromosome seven retained approximately half the autosomal coverage in males, as expected for an × chromosome (Figure 4). Closely related Diptera species are likewise male-heterogametic, supporting an XY sex determination system in BSF. Unlike *Drosophila melanogaster*, the dot-like (Muller F-element) chromosome of BSF is not a redundant sex chromosome and will provide an interesting candidate for chromosome evolution studies [74].

**Figure 4.**
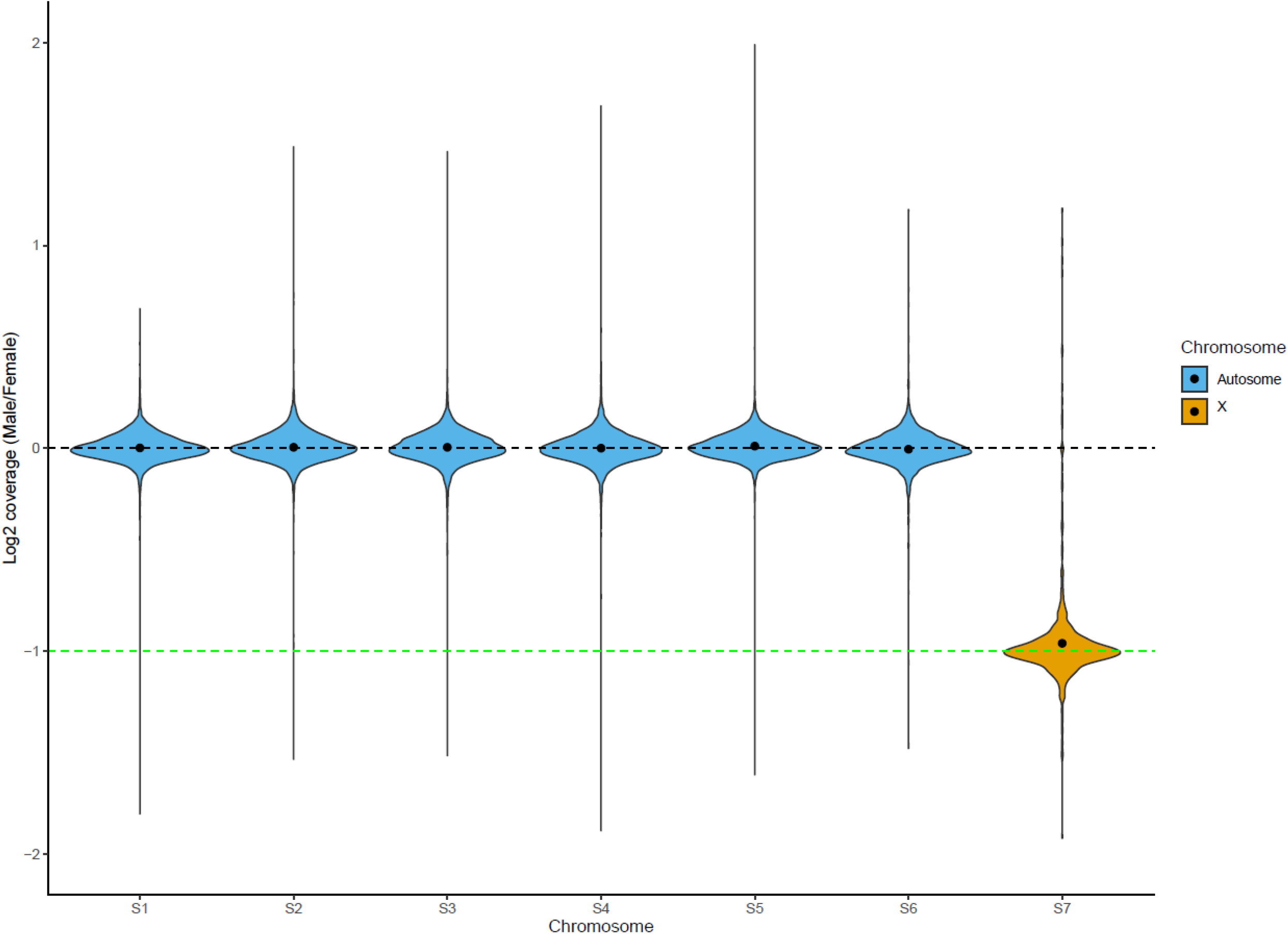
Sex chromosome identification of the *Hermetia illucens* assembly. Log_2_ coverage (male/female) of mapped male (n=5) and female (n=7) whole-genome BGI re-sequencing data. Chromosome seven reveals half the coverage expected of an autosome resembling that of an X sex chromosome in the *H. illucens* assembly.

## Conclusions

We used a combination of PacBio, 10X Genomics linked-reads and Hi-C analysis to assemble the first chromosome-scale BSF genome. A final genome size of 1.01 Gb was produced using 10X and Hi-C scaffolding to obtain contig and scaffold N50 values of 16.01 Mb and 180.46 Mb respectively. We annotated 17,664 protein coding genes using the BRAKER2 pipeline. This chromosome-level assembly provides a significant increase (>100-fold contiguity) in reference quality compared to the existing reference genome. Genomic characterisation identified a sex chromosome and potential candidate regions for further genomic investigations. We also identify the inbreeding coefficient of the established reference population, providing a potential method for inbreeding monitoring in commercial mass rearing BSF facilities. Availability of this novel chromosomal Stratiomyidae reference assembly will aid further research to characterise the genetic architecture behind the unique phenotypic and commercially valuable traits of the BSF. The provided tools will also be of benefit for developing biotechnology resources for genetic manipulation to improve the efficient application of this farmed insect.

## Availability of supporting data

The data sets supporting the results of this article are available under the Bioprojects PRJEB23696 and PRJEB37575.

## Additional files

**Supplementary Figure 1**. GenomeScope profile of *Hermetia illucens* genome using surveyed adults.

**Supplementary Figure 2.** BlobToolKit *Hermetia illucens* GC-coverage by taxonomy.

**Supplementary Figure 3.** Cumulative scaffold length comparing *Hermetia illucens* assembly contiguity.

**Supplementary Figure 4.** Whole genome alignment of *Hermetia illucens* GCA_009835165.1 and iHerIll assemblies.

**Supplementary Figure 5.** Genome wide diversity, Tajima D and ROH analysis of the sampled *Hermetia illucens* EVE population.

**Supplementary Figure 6.** Genome wide diversity, Tajima D and ROH analysis of chromosome five of the *Hermetia illucens* EVE population.

**Supplementary Table 1.** Raw sequencing statistics of *Hermetia illucens* de novo DNA sequencing.

**Supplementary Table 2**. GenomeScope estimated genome characteristics for *Hermetia illucens*.

**Supplementary Table 3.** Genome contiguity statistics of the *Hermetia illucens* genome.

**Supplementary Table 4.** Genome assembly statistics of *Hermetia illucens*.

**Supplementary Table 5.** Full BUSCO table for assembled *Hermetia illucens* and Diptera genome assemblies.

**Supplementary Table 6.** Quality assessment of *Hermetia illucens* assemblies using QUAST.

**Supplementary Table 7.** Genomic diversity and inbreeding of the sampled *Hermetia illucens* EVE population.

## DECLARATIONS

### List of abbreviations

BSF: Black Soldier Fly
BSFL: Black Soldier Fly Larvae
PacBio: Pacific Biosciences
BUSCO: Benchmarking Universal Single-Copy Orthologs
GC: Guaninecytosine
Gb: gigabase
Mb: megabase
kb: kilobase
bp: base pairs
RNA-seq: RNA-sequencing
Hi-C: high-throughput chromosome conformation capture
BWA: Burrows-Wheeler Aligner
PE: paired-end
SMRT: single-molecule realtime
SRA: Sequence Read Archive
LINE: long interspersed nuclear elements
SINE: Short Interspersed Nuclear Elements
LTR: long terminal repeat
STAR: Spliced Transcripts Alignment to a Reference
ROH: Runs Of Homozygosity
SNP: Single Nucleotide Polymorphism.

### Consent for publication

Not applicable.

### Ethics approval and consent to participate

Not applicable.

### Authors contributions

C.D.J, M.P and R.D conceived and supervised the study. M.Q provided genome sequencing services. T.N.G and I.A.W performed DNA extractions. S.A.M performed genome assembly. J.T and Y.S performed genome decontamination. J.M.D.W and T.N.G curated the final genome. T.N.G carried out genome annotation and analysis. T.N.G drafted the manuscript. All authors commented on the manuscript. All authors approve of the manuscript.

## Acknowledgements

We thank the Wellcome Sanger Institute (Cambridge, UK) DNA pipelines team for performing all PacBio, 10X linked-read and Hi-C library preparation and sequencing. Gratitude to Richard Challis for advice with BlobToolKit installation. Also extend thanks to BGI for re-sequencing additional BSF individuals.

