## Supplemental material for "A high-quality, chromosome-level genome assembly of the Black Soldier Fly (*Hermetia Illucens* L.)"

### Supplementary material

**Supplementary Table 1. Raw statistics of *Hermetia illucens de novo* DNA sequencing.** Produced data for the sequencing and assembly of the *H. illucens* genome. Estimated depth calculated based on iHerIll final genome size.

| Input individual | Sex | Platform | Reads produced | Estimated depth (X) |
|---|---|---|---|---|
| Reference pupa | NA | PacBio Sequel | 6 million | 141 |
| Reference pupa | NA | 10X | 391 million | 84 |
| Related adult | Female | Illumina | 444 million | 61 |
| Related adult | Male | Illumina | 479 million | 65 |
| Reference sibling pupa | NA | Hi-C | 478 million | 126 |

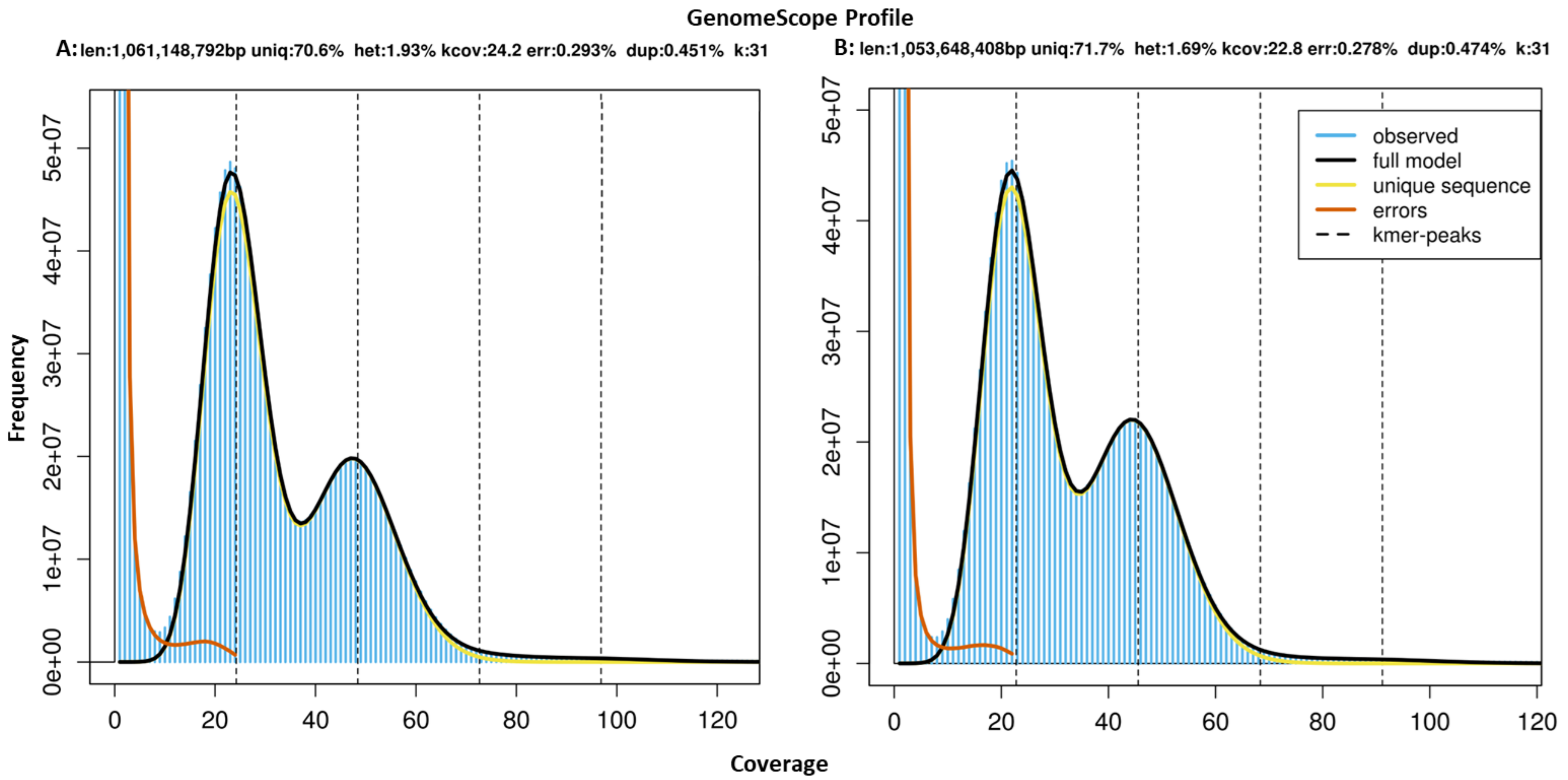


**Supplementary Figure 1. GenomeScope profile of *Hermetia illucens* genome using surveyed adults.** Illumina WGS data of both a male (A) and female (B) *H. illucens* adult. Provisional genome size estimate is 1.06 Gb, repeat content of 46.25% and predicted mean heterozygosity of 1.8%.

**Supplementary Table 2. GenomeScope estimated genome characteristics for *Hermetia illucens.***

| Estimated genome statistic | *Hermetia illucens* | |
|---|---|---|
| | **Female sample** | **Male sample** |
| k-mer size | 31-mer | 31-mer |
| Total genome size (bp) | 1,053,648,408 | 1,061,148,792 |
| Ploidy | Diploid | Diploid |
| Heterozygosity (%) | 1.69 | 1.93 |
| Duplicated sequence content (%) | 45.1 | 47.4 |

**Supplementary Table 3. Genome contiguity statistics of the *Hermetia illucens* genome.**

| Statistics | bp | # |
|---|---|---|
| Sum | 1,004,932,590 | 20 |
| N50 | 180,357,958 | 3 |
| N60 | 173,077,783 | 4 |
| N70 | 173,077,783 | 4 |
| N80 | 116,824,456 | 5 |
| N90 | 103,444,952 | 6 |
| N100 | 11,090 | 20 |
| Largest | 222,122,703 | 1 |

**Supplementary Table 4. Genome assembly statistics of *Hermetia illucens*.** Genome assembly statistics and GC (%) content.

| Scaffold ID | Size (bp) | % | GC (%) |
|---|---|---|---|
| S1 | 222,122,703 | 22.103 | 42.21 |
| S2 | 191,142,057 | 19.020 | 42.41 |
| S3 | 180,357,958 | 17.947 | 42.50 |
| S4 | 173,077,783 | 17.223 | 42.53 |
| S5 | 116,824,456 | 11.625 | 42.21 |
| S6 | 103,444,952 | 10.294 | 43.11 |
| S7 | 15,435,017 | 1.536 | 43.14 |
| scaffold_43_arrow_ctg1 | 900,911 | 0.090 | 40.57 |
| scaffold_49_arrow_ctg1 | 489,531 | 0.049 | 42.74 |
| scaffold_54_arrow_ctg1 | 278,815 | 0.028 | 42.59 |
| scaffold_57_arrow_ctg1 | 166,227 | 0.017 | 45.21 |
| scaffold_58_arrow_ctg1 | 152,185 | 0.015 | 59.00 |
| scaffold_59_arrow_ctg1 | 147,520 | 0.015 | 37.75 |
| scaffold_60_arrow_ctg1 | 90,749 | 0.009 | 45.61 |
| scaffold_61_arrow_ctg1 | 84,925 | 0.008 | 43.54 |
| scaffold_62_arrow_ctg1 | 68,247 | 0.007 | 36.93 |
| scaffold_64_arrow_ctg1 | 63,031 | 0.006 | 42.66 |
| scaffold_65_arrow_ctg1 | 58,701 | 0.006 | 27.20 |
| scaffold_75_arrow_ctg1 | 15,732 | 0.002 | 46.24 |
| scaffold_76_arrow_ctg1 | 11,090 | 0.001 | 41.22 |
| **Genome total** | **1,004,932,590** | **100** | **42.46** |

**Supplementary Table 5. Full BUSCO table for assembled *Hermetia illucens* and Diptera genome assemblies.** BUSCO 'insecta_odb9' and 'diptera_odb9' core gene sets used for analysis. * denotes the reported assembly iHerIll of this manuscript. Reference assemblies obtained from NCBI genome database.

| | BUSCO Insecta | | | | | | | | | | BUSCO Diptera | | | | | | | | | |
|---|---|---|---|---|---|---|---|---|---|---|---|---|---|---|---|---|---|---|---|---|
| **Species** | **Complete** | | **Single-copy** | | **Duplicated** | | **Fragmented** | | **Missing** | | **Complete** | | **Single-copy** | | **Duplicated** | | **Fragmented** | | **Missing** | |
| | **%** | **n** | **%** | **n** | **%** | **n** | **%** | **n** | **%** | **n** | **%** | **n** | **%** | **n** | **%** | **n** | **%** | **n** | **%** | **n** |
| *Hermetia illucens** | 98.60 | 1635 | 97.80 | 1622 | 0.80 | 13 | 0.50 | 8 | 0.90 | 15 | 92.60 | 2592 | 92.2 | 2581 | 0.40 | 11 | 4.0 | 112 | 3.40 | 95 |
| *Hermetia illucens* (GCA_009835165.1) | 98.90 | 1640 | 91.10 | 1510 | 7.80 | 129 | 0.60 | 10 | 0.50 | 8 | 93.60 | 2620 | 87 | 2435 | 6.60 | 185 | 3.7 | 104 | 2.70 | 76 |
| *Drosophila melanogaster* ( GCA_000001215.4) | 99.70 | 1653 | 99.00 | 1641 | 0.70 | 12 | 0.20 | 3 | 0.10 | 2 | 98.7 | 2762 | 98.2 | 2748 | 0.5 | 14 | 0.8 | 21 | 0.5 | 16 |
| *Drosophila virilis* (GCA_000005245.1) | 99.10 | 1643 | 98.10 | 1626 | 1.00 | 17 | 0.40 | 7 | 0.50 | 8 | 98.4 | 2754 | 98.0 | 2743 | 0.4 | 11 | 0.9 | 24 | 0.7 | 21 |
| *Musca domestica* (GCA_000371365.1) | 98.60 | 1635 | 96.90 | 1607 | 1.70 | 28 | 0.40 | 7 | 1.00 | 17 | 97.5 | 2729 | 96.0 | 2686 | 1.5 | 43 | 1.6 | 45 | 0.9 | 25 |
| *Stomoxys calcitrans* (GCA_001015335.1) | 98.40 | 1631 | 97.70 | 1620 | 0.70 | 12 | 1.00 | 17 | 0.60 | 10 | 96.7 | 2706 | 96.2 | 2692 | 0.5 | 14 | 2.1 | 58 | 1.2 | 35 |
| *Glossina morsitans morsitans* (GCA_001077435.1) | 98.90 | 1640 | 96.60 | 1602 | 2.30 | 38 | 0.60 | 10 | 0.50 | 8 | 96.5 | 2703 | 95.7 | 2680 | 0.8 | 23 | 2.4 | 67 | 1.1 | 29 |
| *Anopheles gambiae* (GCA_001542645.1) | 89.40 | 1482 | 85.70 | 1421 | 3.70 | 61 | 7.40 | 123 | 3.20 | 53 | 71.6 | 2002 | 69.7 | 1950 | 1.9 | 52 | 16.0 | 449 | 12.4 | 348 |
| *Aedes aegypti* (GCA_002204515.1) | 98.90 | 1640 | 94.50 | 1567 | 4.40 | 73 | 0.40 | 7 | 0.70 | 12 | 96.7 | 2706 | 92.9 | 2601 | 3.8 | 105 | 1.8 | 50 | 1.5 | 43 |
| *Culex quinquefasciatus* (GCA_000209185.1) | 96.70 | 1603 | 91.80 | 1522 | 4.90 | 81 | 0.80 | 13 | 2.50 | 41 | 93.9 | 2626 | 91.0 | 2546 | 2.9 | 80 | 3.2 | 90 | 2.0 | 83 |

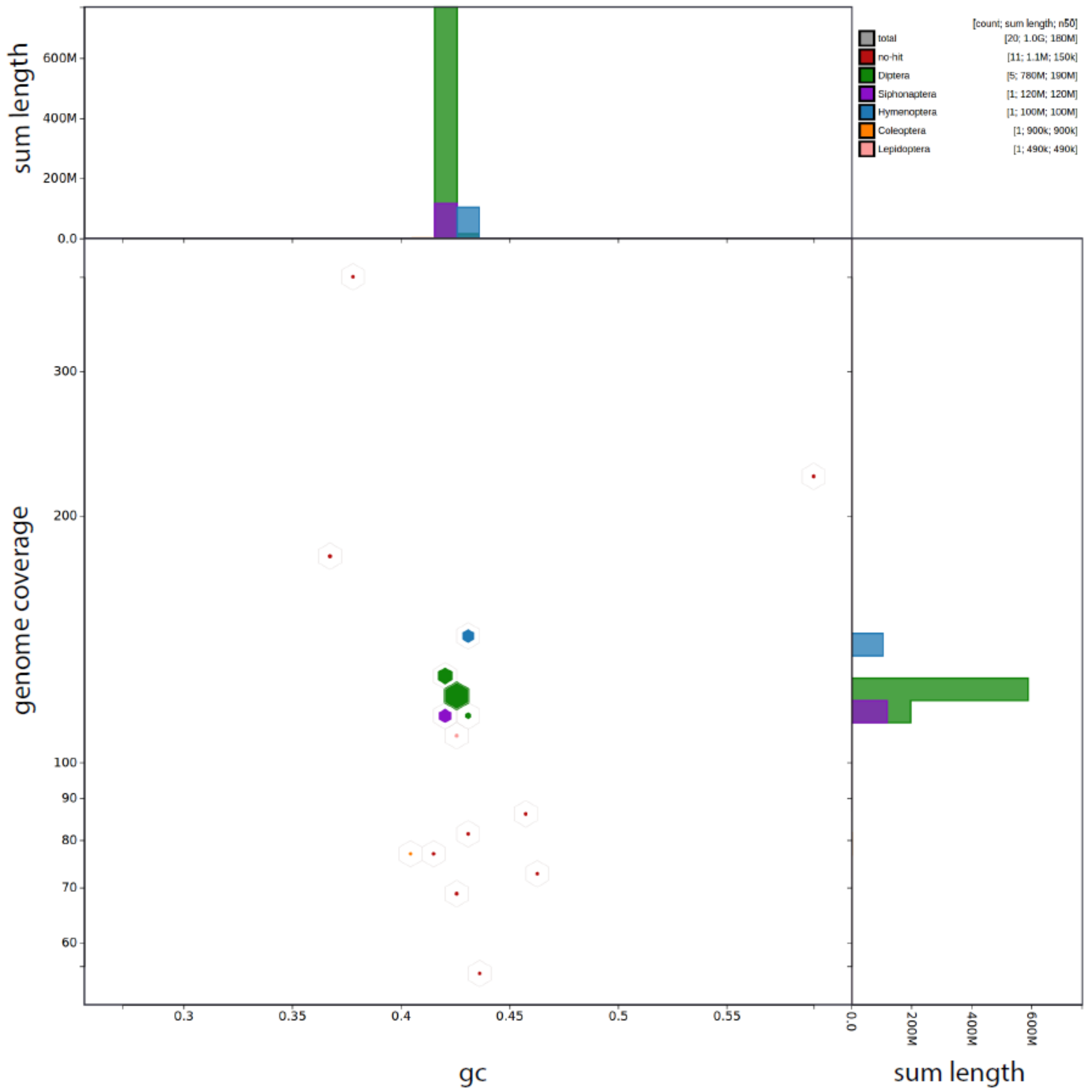


**Supplementary Figure 2. BlobToolKit *Hermetia illucens* GC-coverage by taxonomy.** Taxonomic identification of the assembled Pacific Bioscience sequences used for the primary assembly of the *Hermetia illucens* genome.

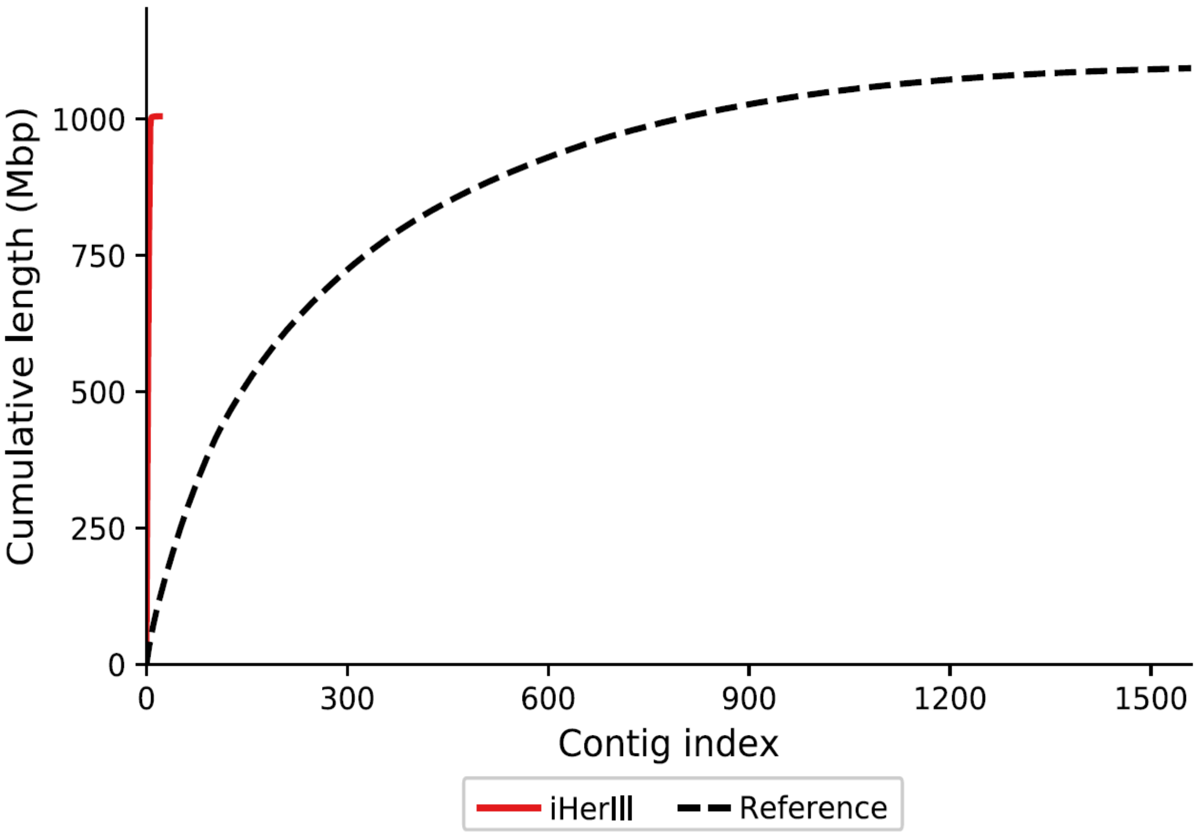


**Supplementary Figure 3. Cumulative scaffold length comparing *Hermetia illucens* assembly contiguity.** A contiguous assembly is observed in the iHerIll assembly when compared to GCA_009835165.1 as a reference assembly. Contiguity represented by a vertical line and minimal horizontal tail showing cumulative length is contained within a minimum contig index of the iHerIll assembly.

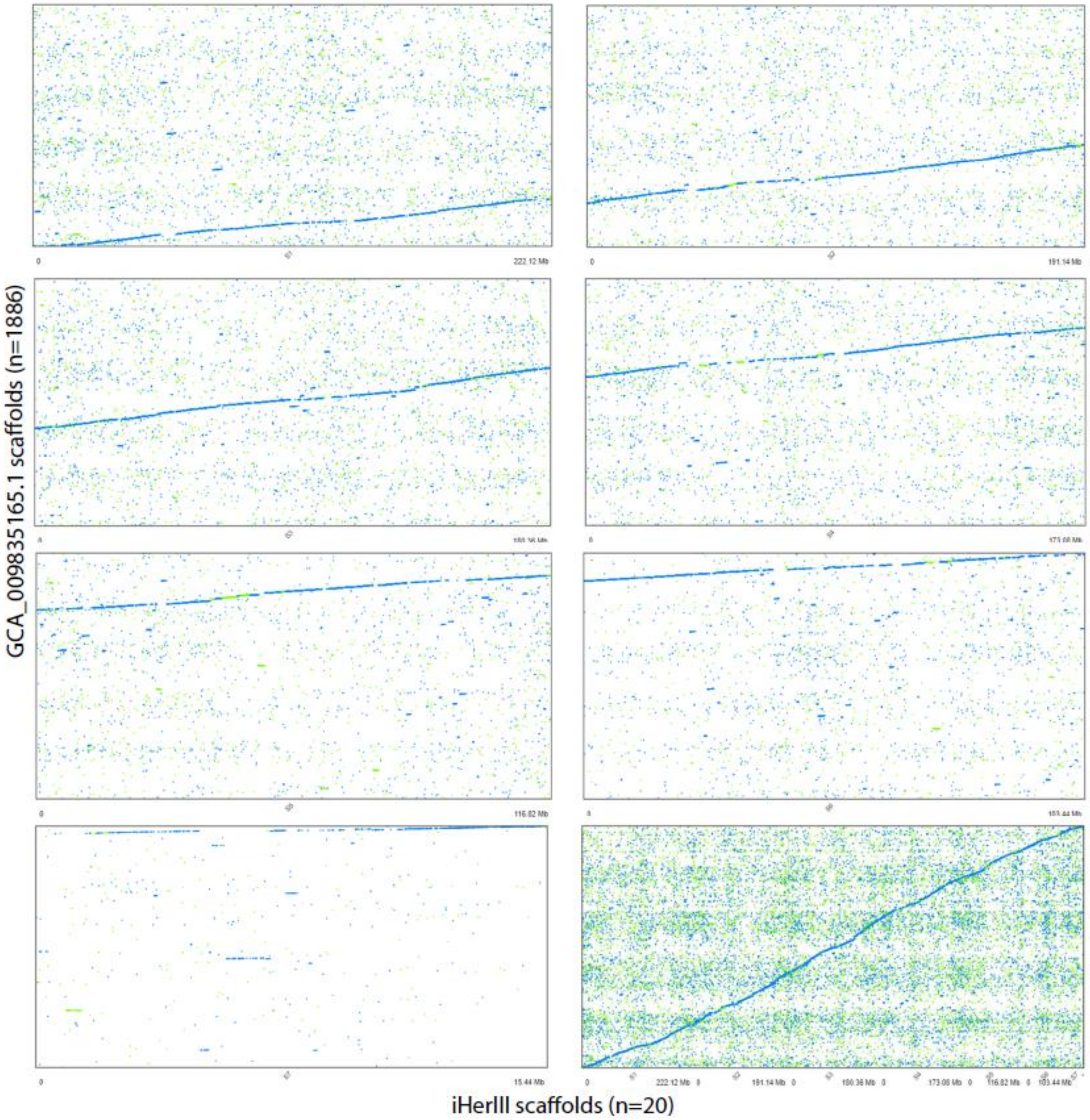


**Supplementary Figure 4. Whole genome alignment of *Hermetia illucens* GCA_009835165.1 and iHerIll assemblies.** Chromosome alignment of GCA_009835165.1 scaffolds to the iHerIll assembly both per chromosome (top left to bottom left) and whole genome overview (bottom right). Unique forward alignments = blue, unique reverse alignments = orange and repetitive sequences = green.

**Supplementary Table 6. Quality assessment of *Hermetia illucens* assemblies using QUAST.** Quality assessment of GCA_009835165.1 (query) using iHerIll as the reference assembly. Alignment statistics, NA50 and LA50 provide contiguity values based on aligned contigs only.

| **Query statistics** | **Query genome: GCA_009835165.1** |
|---|---|
| Reference genome used | iHerIll |
| Genome size (bp) | 1,099,000,393 |
| Largest alignment (bp) | 3,709,431 |
| NA50 (bp) | 48,152 |
| LA50 (bp) | 2,595 |
| **Estimated misassembly events (#)** | **10,773** |
| *Relocations (#)* | *3,482* |
| *Translocations (#)* | *7,157* |
| *Inversions (#)* | *134* |
| Estimated misassembled contigs *(#)* | 787 |
| Fully unaligned contigs *(#)* | 10 |
| Fully unaligned non-repetitive sequence length (bp) | 302,401 |

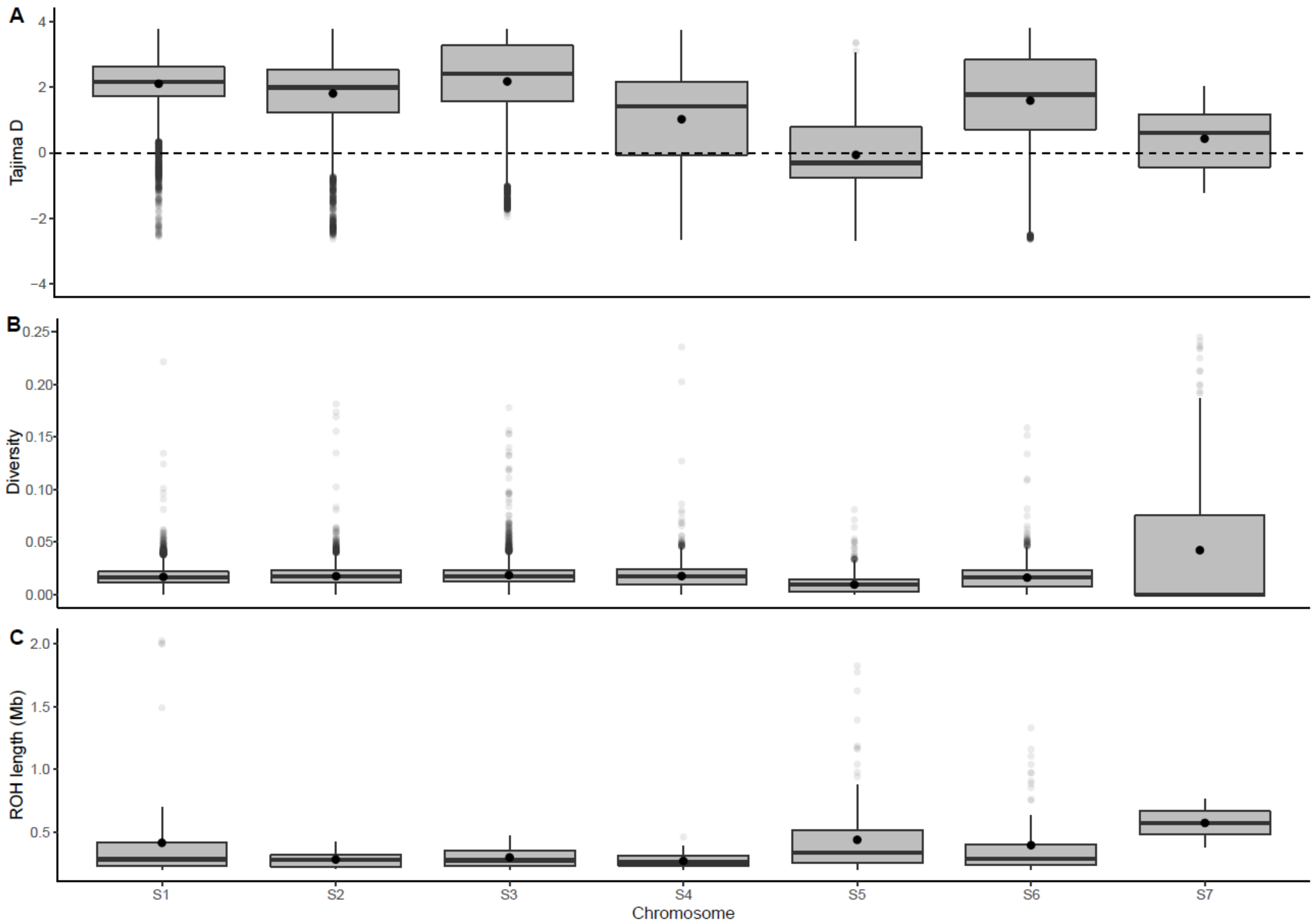


**Supplementary Figure 5. Genome wide diversity, Tajima D and ROH analysis of the sampled *Hermetia illucens* EVE population.** Tajima D (A), nucleotide diversity (B) and Runs of Homozygosity (ROH) values for each of the seven assembled chromosomes. Contigs < 1 Mb (n=13) not included.

**Supplementary Table 7. Genomic diversity and inbreeding of the sampled *Hermetia illucens* EVE population.** Statistics describing mean *H. illucens* sequence diversity, Tajima D values, ROH counts, ROH mean lengths and inbreeding coefficients of the sampled population. Reported statistics generated on a per chromosome basis and genome wide. [a] Final genome wide $F_{ROH}$ excludes identified sex chromosome seven (S7).

| Query sequence | Pi (π) | Tajima D | ROH (#) | ROH mean length (kb) | $F_{ROH}$ |
|---|---|---|---|---|---|
| S1 | 0.017 | 2.1 | 68 | 415.43 | 0.011 |
| S2 | 0.018 | 1.802 | 37 | 282.13 | 0.005 |
| S3 | 0.019 | 2.170 | 21 | 299.14 | 0.003 |
| S4 | 0.018 | 1.019 | 38 | 270.68 | 0.005 |
| S5 | 0.010 | -0.066 | 193 | 439.10 | 0.060 |
| S6 | 0.016 | 1.586 | 85 | 396.49 | 0.027 |
| S7 | 0.072 | 0.428 | 2 | 573.86 | 0.040 |
| Genome Wide | 0.017 | 1.576 | 444 | 393.81 | 0.019 |

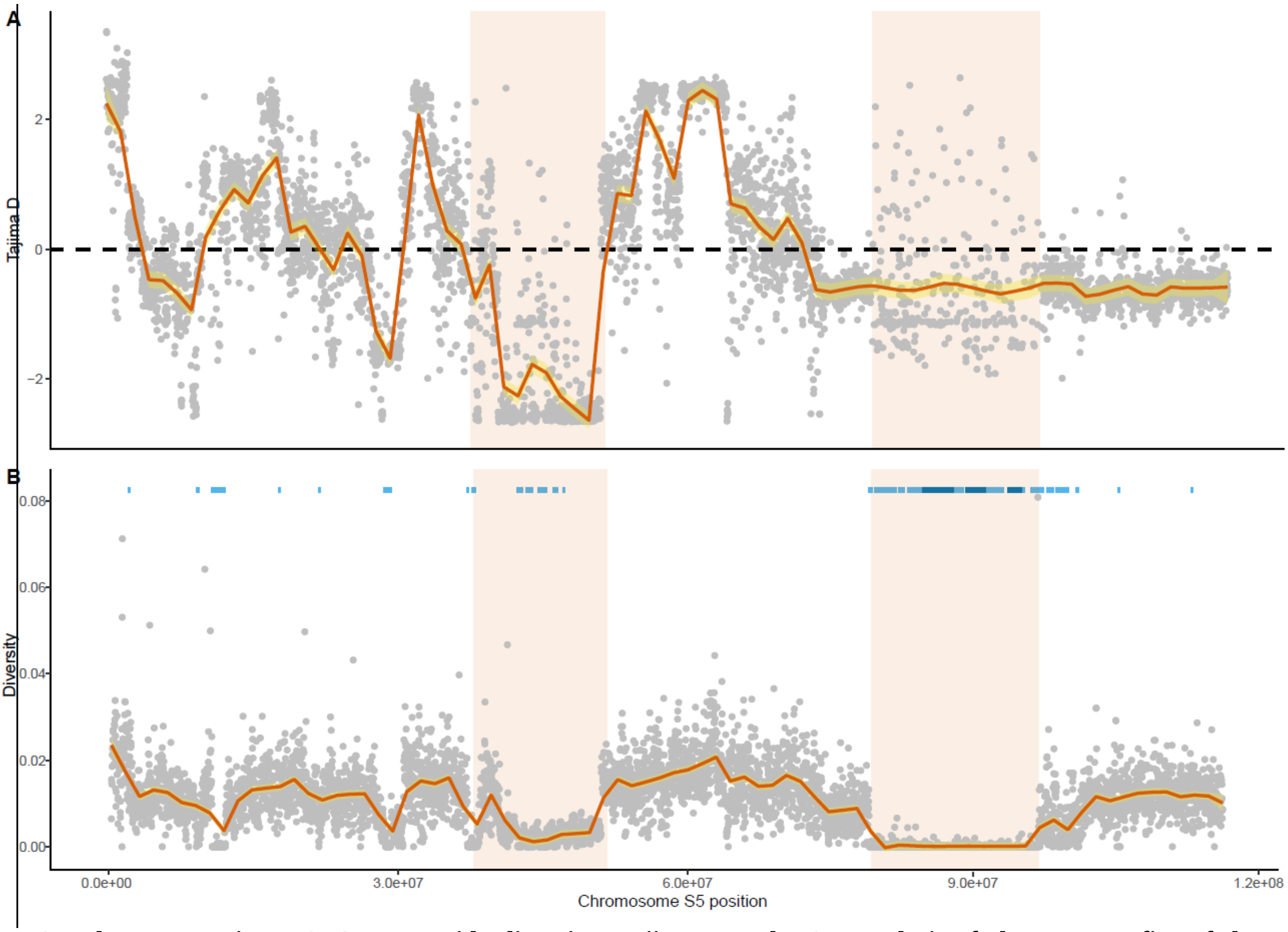


**Supplementary Figure 6. Genome wide diversity, Tajima D and ROH analysis of chromosome five of the *Hermetia illucens* EVE population.** Tajima D (A), nucleotide diversity and Runs of Homozygosity (ROH) (B) values across genomic positions of chromosome five. Mean 20 kb windows smoothed with local regression and regions of interest highlighted. Both short (light blue) and long (dark blue) ROH shown across genome position.
